# Chromosome-level *de novo* assembly of *Coprinopsis cinerea A43mut B43mut pab1-1* #326 and genetic variant identification of mutants using Nanopore MinION sequencing

**DOI:** 10.1101/2020.11.09.367581

**Authors:** Yichun Xie, Yiyi Zhong, Jinhui Chang, Hoi Shan Kwan

## Abstract

The homokaryotic *Coprinopsis cinerea* strain *A43mut B43mut pab1-1* #326 is a widely used experimental model for developmental studies in mushroom-forming fungi. It can grow on defined artificial media and complete the whole lifecycle within two weeks. The mutations in mating type factors *A* and *B* result in the special feature of clamp formation and fruiting without mating. This feature allows investigations and manipulations with a homokaryotic genetic background. Current genome assembly of strain #326 was based on short-read sequencing data and was highly fragmented, leading to the bias in gene annotation and downstream analyses. Here, we report a chromosome-level genome assembly of strain #326. Oxford Nanopore Technology (ONT) MinION sequencing was used to get long reads. Illumina short reads was used to polish the sequences. A combined assembly yield 13 chromosomes and a mitochondrial genome as individual scaffolds. The assembly has 15,250 annotated genes with a high synteny with the *C. cinerea* strain Okayama-7 #130. This assembly has great improvement on contiguity and annotations. It is a suitable reference for further genomic studies, especially for the genetic, genomic and transcriptomic analyses in ONT long reads. Single nucleotide variants and structural variants in six mutagenized and cisplatin-screened mutants could be identified and validated. A 66 bp deletion in Ras GTPase-activating protein *(RasGAP)* was found in all mutants. To make a better use of ONT sequencing platform, we modified a high-molecular-weight genomic DNA isolation protocol based on magnetic beads for filamentous fungi. This study showed the use of MinION to construct a fungal reference genome and to perform downstream studies in an individual laboratory. An experimental workflow was proposed, from DNA isolation and whole genome sequencing, to genome assembly and variant calling. Our results provided solutions and parameters for fungal genomic analysis on MinION sequencing platform.

**Highlight:**

- A chromosome-level genome assembly of *C. cinerea #326*
- A fast and efficient high-molecular-weight fungal genomic DNA isolation protocol
- Structural variant and single nucleotide variant calling using Nanopore reads
- A series of solutions and reference parameters for fungal genomic analysis on MinION

## 1. Introduction

Genomic data are powerful resources to understand the complex mechanisms and biological processes. They support the genetics, genomics and phylogenetics studies in wide ranges and fine scales (Kono and Arakawa, 2019). High-quality reference genomes are critical for the downstream analyses, such as gene detection, functional annotation, variant determination and evolutionary analysis (Xu et al., 2016). Building a *de novo* genome assembly is one of the fundamental steps in genome analysis of novel sequences.

Hybrid genome assembly methods construct the genome using error-prone long reads and accurate short reads (Antipov et al., 2016; Díaz-Viraqué et al., 2019; Miller et al., 2017; Wick et al., 2017). *De novo* assemblies using Oxford Nanopore Technologies (ONT) long read sequences have the ease of constructing the chromosomes in single contigs, with telomere repeats on chromosome flanks (Kolmogorov et al., 2018; Salazar et al., 2017). Moreover, long reads can span the entire tandems of repeats in the genome, resolve the complex regions and improve contiguity that short reads could not achieved (Shin et al., 2019; Tyson et al., 2018). On the other hand, short read polishing could make up the problem of high error rate in long read assembly and cut down the requirement of sequencing depth (Wang and Au, 2020). Therefore, hybrid assembly method greatly improves the genome assembly quality and cuts down the unit price of data generation (Díaz-Viraqué et al., 2019; Miller et al., 2017; Tan et al., 2018).

High-molecular-weight genomic DNA (HMW gDNA) samples are needed for genome sequencing and downstream analyses. In Nanopore experiments, sequencing read length greatly influences the contiguity of the final assembly (Goldstein et al., 2019), and it is mostly restricted by the length of DNA molecules and the ability of delivering HMW DNA molecules to the sequencing pores (Laver et al., 2015). Therefore, isolating HMW gDNA from samples would be the start point toward a successful long read sequencing and genome assembly. Generally, commercial spin column kits can extract the genomic DNA of most samples within one hour, yielding 3 ~ 8 μg DNA/100 mg fresh mycelia. However, the length distribution of DNA fragment was not ideal, and those sequencing data might not be sufficient for chromosome-level genome assembly. Although several HMW DNA extraction kits are now available on the market, most of them are designed for animal samples or bacteria, thus, extra reagents and manipulations are required to make them suitable for plant and fungal samples. Classical DNA extraction methods, such as CTAB method and phenolchloroform method, are applicable to isolate 100-200 Kbp DNA fragment from most biological samples, but these methods have the drawback of intensive labour and hazardous reagent exposure (Gong et al., 2019; Schwessinger and Rathjen, 2017; Vaillancourt and Buell, 2019). Further, plug extraction is the best method to acquire large DNA fragments in megabase-sized (Quick and Loman, 2018). Such ultra-long DNA fragments are good for gigabase-sized genome assembly, genome mapping and bacterial artificial chromosome (BAC) library construction (Chaney et al., 2016; Hu et al., 2019; H.-B. Zhang et al., 2012). However, plug extraction method generally takes three days to obtain the final product, and requires good experimental skills and special laboratory setup (M. Zhang et al., 2012). Magnetic beads have long been used to purified DNA in molecular biology studies (Berensmeier, 2006). Library preparation protocols distributed by ONT, PacBio and Illumina all include the sample DNA and library purification steps using magnetic beads (e.g. AMPure XP beads, Beckman Coulter, USA). However, direct DNA isolation and purification from cell lysis using magnetic beads has not been widely tested and further modifications can be applied (He et al., 2017; Mayjonade et al., 2017).

*Coprinopsis cinerea* is one of the model organisms that commonly used in fungal studies. It is a typical mushroom-forming fungus with several developmental destinies (Kües, 2000). The karyotype and genetic features of *C. cinerea* have been well studied. It is known that *C. cinerea* generally has 13 chromosomes and the estimated genome size of 37.5 Mbp (Holm et al., 1981; Stajich et al., 2010). To date, genome resources of two *C. cinerea* strains have been released. The genome of the monokaryotic strain Okayama-7 #130 was sequenced using whole genome shotgun sequencing method and constructed to the chromosome-level (Stajich et al., 2010). Another strain, the homokaryotic strain *A43mut B43mut pab1-1* #326, is a self-fertilise strain which can fruit without mating. It has widely been used in developmental studies (Lau et al., 2020; Masuda et al., 2016; Muraguchi et al., 2015; Sakamoto et al., 2017; Sugano et al., 2017). The genome of strain #326 was sequenced using Illumina HiSeq 2000 platform and assembled into 944 scaffolds (Muraguchi et al., 2015). When applying the assembly to long read data analysis, the fragmented genome sequence raises difficulties to its application, such as genetic variant calling and genomic region determination. Therefore, a better genome sequence of strain #326 is in need.

Here, we take *C. cinerea A43mut B43mut pab1-1* #326 as an example, records a working procedure for fungal genomic data collection and processing. The workflow includes HMW genomic DNA isolation, genome sequencing, genome assembly and annotation, as well as genetic variant calling of relative strains. We aim to (i) obtain an efficient HMW gDNA isolation protocol that is suitable for fungal samples; (ii) achieve a chromosome-level reference genome for *C. cinerea* strain #326 using the hybridisation of long-read and shortread sequencing technologies; (iii) perform genome annotation and comparison on long read and short read assemblies; (iv) apply the assembly in structural variant and single nucleotide variant identification of mutated fungal isolates. Our results shall provide a series of experimental and *in silico* references on fungal genomic analyses using MinION sequencing platform.

## 2. Materials and methods

### 2.1 Strains and cultivation

The homokaryotic fruiting strain *C. cinerea A43mut B43mut pab1-1* #326 was grown at 37°C on solid YMG medium (0.4% yeast extract, 1% malt extract, 0.4% glucose and 1.5% agar, 36 g medium per 90 mm diameter petri dish, same below). Fresh vegetative mycelia were collected on day 5, and polysaccharide-rich sclerotia-forming mycelia were collected on day 12. In addition, three fruiting and three non-fruiting mutated isolates were included in this study. Details of mutagenesis and isolation were described in Text S1. DNA sample of mutants were extracted using DNeasy Plant Mini Kit (Qiagen, Germany). The workflow of data collection and processing was summarised in Fig. 1.

**Fig. 1.**
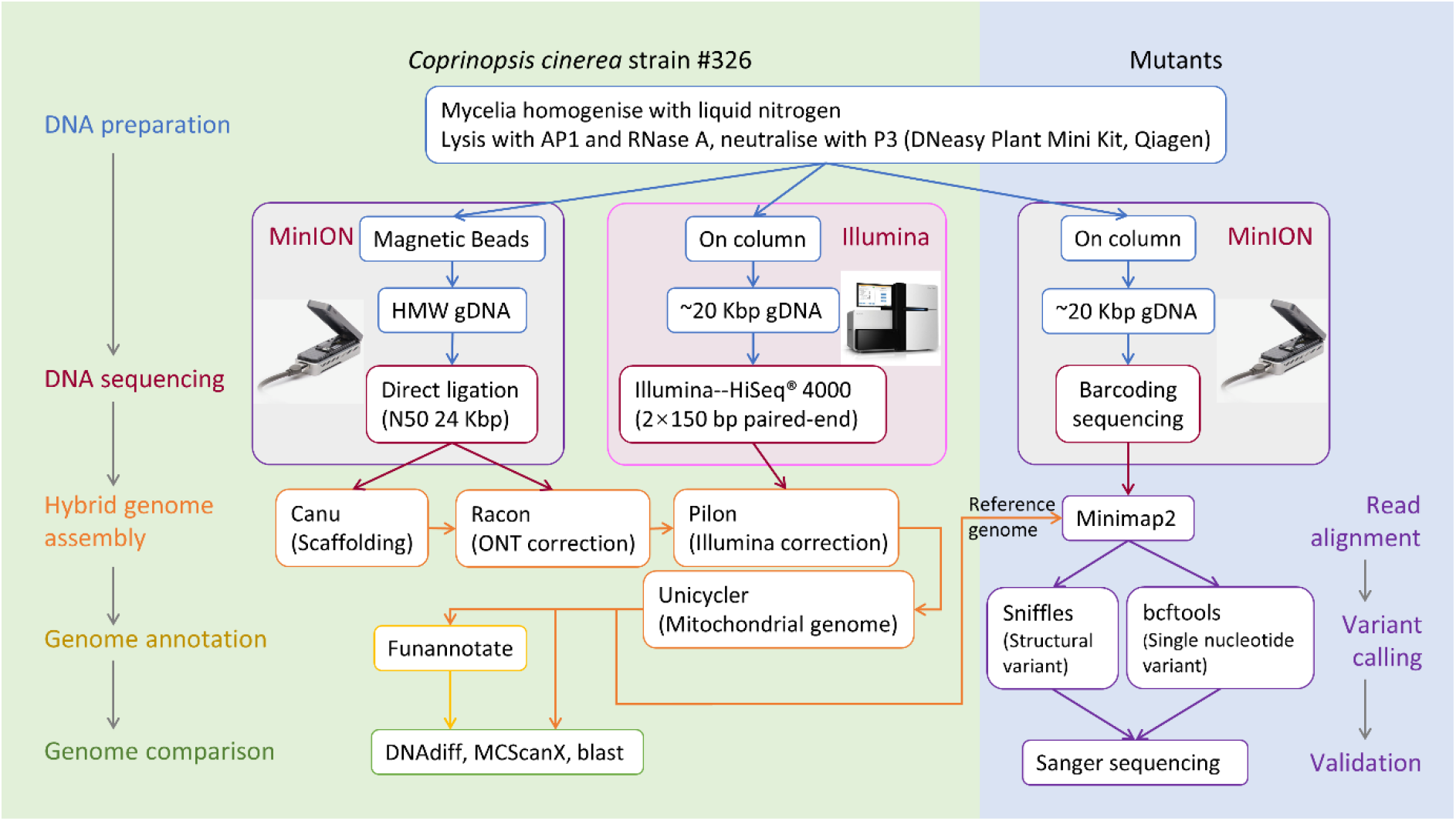
Flowchart for reference genome construction and genetic variant calling of mutants.

### 2.2 High-molecular-weight genomic DNA extraction

A step-to-step HMW gDNA isolation protocol was described in detail in Text S2. In short, fresh mycelia were homogenised using liquid N_2_. Cell lysis and neutralisation were performed using buffer AP1, RNase A and buffer P3 in DNeasy Plant Mini Kit (Qiagen, Germany). DNA purification was done using AMPure XP beads (Beckman Coulter, USA) with modifications (Mayjonade et al., 2017). The DNA binding step was repeated to increase the total yield. DNA quality and quantity were determined using Nanodrop (Thermofisher, USA), Qubit^®^ dsDNA HS Assay Kits and Qubit^®^ 2.0 fluorometer (Invitrogen, USA). The DNA fragment size was estimated using 0.8 % agarose gel electrophoresis.

### 2.3 MinION library preparation, sequencing and basecalling

The Nanopore sequencing library was prepared using ligation sequencing kit SQK-LSK109 and other specific reagents following the genomic DNA by ligation sequencing protocol (for strain #326, version: GDE_9063_v109_revQ_14Aug2019) and PCR barcoding (96) genomic DNA sequencing protocol (for mutants, version: PBGE96_9068_v109_revM_14Aug2019) released by the manufacturer (Oxford Nanopore Technologies, England). Common modifications were as follows: (i) Regarding the high-molecular-weight of the prepared DNA, the amount of input DNA was increased from ~ 1 μg to ~ 1.5 μg; (ii) DNA repair and end-prep cocktail was incubated at 20°C for 30 min and 65°C for 30 min in a thermal cycler; (iii) For direct adapter ligation of strain #326 DNA, adapter ligation cocktail was incubation at 16°C for 12 h; for PCR barcoding adapter ligation of DNA from mutated isolates, adapter ligation cocktail was incubation at 20°C for 1 h; (iv) DNA was eluted by incubating at 50°C for 10 min. Qubit fluorometer was used to quantify DNA content in eluted solution. DNA recovery rate was also calculated.

Library priming and loading was done following the manufacturer’s instruction. Sequencing of strain #326 and mutant strains were run on separate MinION flow cell FLO-MINSP6 R 9.4.1 (Oxford Nanopore Technologies, England). Both flow cells ran for 72 h under the control of MinKNOW v3.6.0. For whole genome sequencing of strain #326, 250 μl priming mix was added from priming port to refuel the flowcell after running for 6 h and extra 6 μl DNA library was loaded to the flowcell after sequencing for 8 h and 20 h.

High accuracy base calling and filtering were done by “guppy_basecaller” in Guppy v3.4.3 GPU released by Oxford Nanopore Technologies (https://community.nanoporetech.com). The default parameters, including q score cut-off of 7 (accuracy rate over 85 %) and config file “dna_r9.4.l_450bps_hac.cfg”, were used in base calling. Barcodes were identified by “guppy_barcoder” in Guppy. Quality control of reads was performed using an R base package MinIONQC v1.3.5 (Lanfear et al., 2019; R Core Team, 2020).

### 2.4 Illumina library preparation, sequencing and quality control

*C. cinerea* strain #326 genome was also resequenced with Illumina platform. DNA sample was extracted using DNeasy Plant Mini Kit (Qiagen, Germany). DNA sequencing library was prepared with insert size of 270 bp and sequenced with Illumina HiSeq^®^ 4000 at the 2×150 bp paired-end read mode by the Beijing Genomics Institute (BGI, Shenzhen, China). Adapter trimming and quality filtering of DNA sequences was performed using fastp (Chen et al., 2018) with default parameters (window size of 4, mean quality of Q 20, and minimum read length of 15 nt).

### 2.5 *De novo* genome assembly using Nanopore and Illumina reads

Hybrid genome assembly was performed following the ONT assembly and Illumina polishing pipeline (https://github.com/nanoporetech/ont-assembly-polish, Oxford Nanopore Technologies, England). Canu v2.0 was used to assemble the Nanopore reads (Koren et al., 2017). Genome size of *C. cinerea* #326 was estimated to be 37.5 Mb according to the genome of *C. cinerea* Okayama-7 #130 (Stajich et al., 2010). The parameter “trimReadsCoverage”, which indicates the minimum depth of evidence to retain bases, was increased from one read to five reads to minimise the impact of sequencing mistakes on genome assembly. Other parameters in Canu were set to default. Canu contigs were polished using Racon v1.4.3 (Vaser et al., 2017). To improve the accuracy, quality filtered Illumina reads were introduced to the assembly and polishing pipeline. Correction of the contigs was done using Pilon v1.20 (Walker et al., 2014). Vector and contamination were removed by NCBI genome processing system. Mitochondrial draft scaffold was manually identified and reassembled using Unicycler v0.4.8 (Wick et al., 2017).

Assembly quality was evaluated by QUAST v5.0.2 (Gurevich et al., 2013). Assessment of genome completeness was done in BUSCO v4.0.5 with basidiomycota_db10 database (Seppey et al., 2019; Simão et al., 2015). Genome was also aligned and blasted against the reference genome of strain #130. Differences of two genomes were compared using dnadiff in MUMmer package v4.0 (Marçais et al., 2018). Genome alignment results were visualized using mummerCoordsDotPlotly.R in dotPlotly (https://github.com/tpoorten/dotPlotly) with minor customisation on plotting. Macrosyntheny and collinearity analysis were performed using the python version MCscan in JCVI utility libraries (Tang et al., 2015; Wang et al., 2012). Scaffolds were compared to the chromosomes of strain #130. To compare the sequence differences between two genome assemblies of strain #326, scaffolds of the short read assembly were align to the hybrid assembly using minimap2 with “-x asm5” option (Li, 2018). The distribution of additional sequences in the genome were visualised with ‘ideogram()’ function in R package RIdeogram v0.2.2 (Hao et al., 2020; R Core Team, 2020). To figure out the minimum coverage required for a sufficient chromosome-level assemble, sequencing reads were randomly sampled for three times using “seqtk sample” to achieve the coverage of 30 ×, 60 ×, 90 ×, 100 × and 120 × (Goldstein et al., 2019). The sub-sequence batches undergone the assembly and polishing pipeline with same parameters as the original one.

### 2.6 Gene prediction and genome annotation

Genome annotation of the draft genome was done using funannotate v1.7.2, following the workflow of clean-sort-mask-train-predict-update-annotate (Palmer and Stajich, 2019). A total of 18 transcriptomes was acquired from NCBI BioProject PRJNA573620 (Xie et al., 2020) and used to help training models for gene prediction and annotation. The RNA-seq data were quality filtered using fastp with default parameters, including the window size of 4, mean quality of Q 20 and minimum read length of 15 nt (Chen et al., 2018). Repeat masked assembly was used to run “funannotate train” function with the parameter “--stranded no -- jaccard_clip --no_trimmomatic”, to align quality filtered RNA reads, and to run trinity and PASA. PASA genome models were used as evidences in the following gene prediction steps. Prediction was performed using “funannotate prediction” function with the parameter “-- busco_db basidiomycota --genemark_mode ET --busco_seed_species coprinus_cinereus -- augustus_species coprinus_cinereus --optimize_augustus”. Gene predictions were generated from different sources, including GeneMark: 13,349, HiQ: 7,089, pasa: 8,803, CodingQuarry: 16,821, Augustus: 7,968, total: 54,030. Gene models were pass to Evidence Modeler with the EVM weight of GeneMark: 1, HiQ: 2, pasa: 6, proteins: 1, CodingQuarry: 2, Augustus: 1, transcripts: 1, to generate final gene models. Untranslated regions (UTRs) were added to the annotation with PASA in “funannotate update” function. Protein sequences were blast against non-redundant (nr) version 5 database (https://ftp.ncbi.nlm.nih.gov/blast/db/v5/) using blastp in NCBI-BLAST+ v2.10.0 (https://ftp.ncbi.nlm.nih.gov/blast/executables/blast+/2.10.0/ncbi-blast-2.10.0+-x64-linux.tar.gz) to improve the annotation. Final annotation file was generated using “funannotate annotate” function. Repeat sequences were statistics using RepeatMasker v4.0.9 (Tarailo-Graovac and Chen, 2009).

### 2.7 Identification and validation of genetic variants in mutated strains

Six mutants were sequenced using MinION sequencing platform. The ONT reads were aligned to the genome assembly using minimap2 v2.17 (Li, 2018). Structural variants (SV) were identified following the minimap2-Sniffles pipeline (Cameron et al., 2019) using Sniffles v1.0.11 (Sedlazeck et al., 2018). Each SV required at least 5 supporting reads and mitochondrial genome was excluded. Single nucleotide variants (SNV) were identified using “mpileup” in bcftools v1.8 (Li et al., 2009). Each SNV has at least 5 supporting reads and the ratio of alternative reads is over 90 %.

SVs and SNVs were validated by PCR amplification and Sanger sequencing. Rapid isolation of DNA templates of 100 mutated isolates were done following a microwave-base DNA extraction protocol (described in Text S1) (Dörnte and Kües, 2013). PCR amplification of targeted sequence were performed with KAPA HiFi HotStart ReadyMix PCR kit (Roche, Germany) and the following program: 95°C for 3 min, followed by 30 cycles 98°C for 20 sec, 65°C for 15 sec, and 72°C for 45 sec, and 72°C for 1 min. PCR products were detected on 1.5 % agarose gel. Sanger sequencing on PCR products were performed by BGI (Shenzhen, China).

### 2.8 Gene expression quantification using RT-qPCR

Total RNA of the original #326 (wildtype) and mutants were isolated from vegetative mycelia and hyphal knots using TRI reagent^®^ (Sigma-Aldrich, USA) following the manufacturer’s instruction with three biological replicates. The RNA was treated with TURBO DNA-*free*™ Kit (Invitrogen, USA) to remove the DNA contamination. Quality control of RNA was performed using 1.5 % agarose gel electrophoresis and NanoDrop™. cDNA was synthesised from the total RNA with anchored-oligo(dT)_18_ primer and random hexamer primer using Transcriptor First Strand cDNA Synthesis Kit (Roche, Germany). One μg of RNA was added to each 20 μl RT reaction. The template-primer mix was denaturised at 65°C, and the RT reaction was incubated as follows: 10 min at 25°C, 30 min at 55°C and 5 min at 85°C. Expressions of selected genes were quantified using SsoAdvancedTM Universal SYBR^®^ Green Supermix (Bio-Rad, USA) with Applied BiosystemsTM 7500 fast Real-Time PCR System (Applied Biosystems, USA). PCR reactions were performed as follows: 30 sec at 95°C, followed by 40 cycles of 15 sec at 95°C and 30 sec at 60°C, instrument default setting on melt-curve analysis. 18S rRNA was used as the internal control. Primers used in this study are listed in Table S1.

### 2.9 Data availability

Genome assembly was submitted to NCBI under the BioProject accession number PRJNA573619. Annotation files are listed in dataset S1. Raw sequences of strain #326 with both Nanopore long read sequencing and Illumina short read sequencing were submitted to NCBI SRA (http://www.ncbi.nlm.nih.gov/sra) under BioProject PRJNA573619. Transcriptomes were under BioProject PRJNA573620. ONT reads of mutated isolates were under BioProject accession numbers PRJNA648015.

## 3. Results

### 3.1 A fast and efficient high-molecular-weight gDNA isolation protocol applicable to Nanopore sequencing

The high-molecular-weight (HMW) genomic DNA isolation was performed using the magnetic beads selective binding method (Mayjonade et al., 2017). Single addition of lysis supernatant to the pre-treated beads resulted in the final DNA yield of around 1 μg per reaction tube. No significant difference was observed on yields between vegetative mycelia and polysaccharide-rich (> 60 % dry weight) sclerotia forming mycelia. Repeating the lysis supernatant addition and binding step increased the final yield to over 10 μg per reaction tube. The genomic DNA was size selected and concentrated to ~ 300 ng/μl in 80 μl by pooling three tubes of DNA products together and performing an extra 1 × volume beads purification. The DNA solution were later diluted to the concentration of 40 ng/μl for library preparation. The length of DNA molecules estimated by gel electrophoresis were over 20 Kbp (Fig. 2). To summarise up, the modified DNA extraction protocol could yield at 10 μg genomic DNA with no size selection using ~ 100 mg sample in 2.5 h or 25 μg HMW genomic DNA using ~ 300 mg sample in 3 h.

**Fig. 2.**
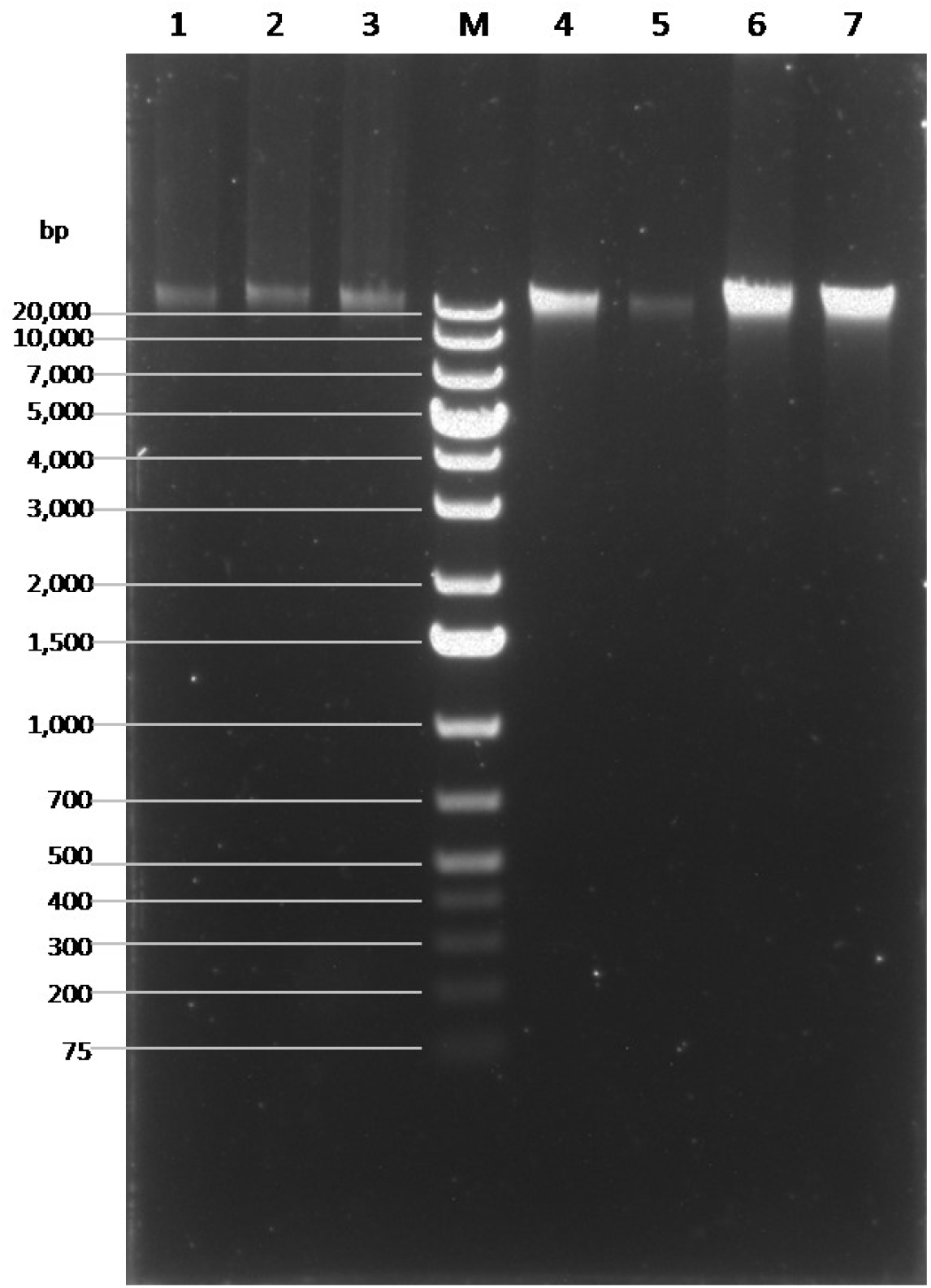
Gel electrophoresis of high-molecular-weight gDNA. 3 μl solution injected to each lane. Condition: 0.8 % agarose gel, 140 V (250 mA) for 30 min. M: GeneRuler 1kb Plus DNA ladder (Thermo Scientific, USA). 1-3: gDNA acquired from a single AMPure tube, lysis supernatant binds to AMPure beads for once; 4-6: gDNA, lysis supernatant binds to AMPure beads for twice. 4: gDNA acquired from a single AMPure tube; 5: 10 times diluted gDNA acquired from a single AMPure tube; 6: gDNA acquired from three AMPure tubes with an extra 1.8 × AMPure beads DNA purification and concentration. 7: DNA library ready for priming and sequencing. Sample 1-2 were isolated from sclerotia forming mycelia (high polysaccharide content); sample 3-7 were isolated from vegetative mycelia.

The HMW DNA library of strain #326 was sequenced on a MinION flowcell with total run time of 72 h. Sequencing read length and Q value were not significantly varied by time (Fig. S1a and b). Due to the loss of sequencing pores, data generation speed significantly dropped down after 40 h of sequencing runtime (Fig. 3a and Fig. S1c). A total of 757,411 reads and 9.44 gigabases were sequenced in 72 h. Mean Q value and median Q value were 10.6 and 11.3. After high accuracy base calling and quality filtering using Guppy, a total of 671,729 reads (88.6 %) and 8,970,263,434 bases (95 %) were acquired. The mean Q value and median Q value were 11.3 and 11.5, representing the mean accuracy of bases at 93 %. The N50 length reached 24,630 bases after filtering and the longest read was 173,441 bases. Over 60 % of bases were acquired from the over 20 kb reads (25.3 %), and 7.3 % of bases were from the over 50 kb reads (1.66 %) (Fig. 3b). In short, most sequencing reads were high quality in accuracy and read length (Fig. 3c-e). These results also proved that our DNA isolation protocol can produce high quality long DNA fragments that are sufficient for Nanopore sequencing.

**Fig. 3.**
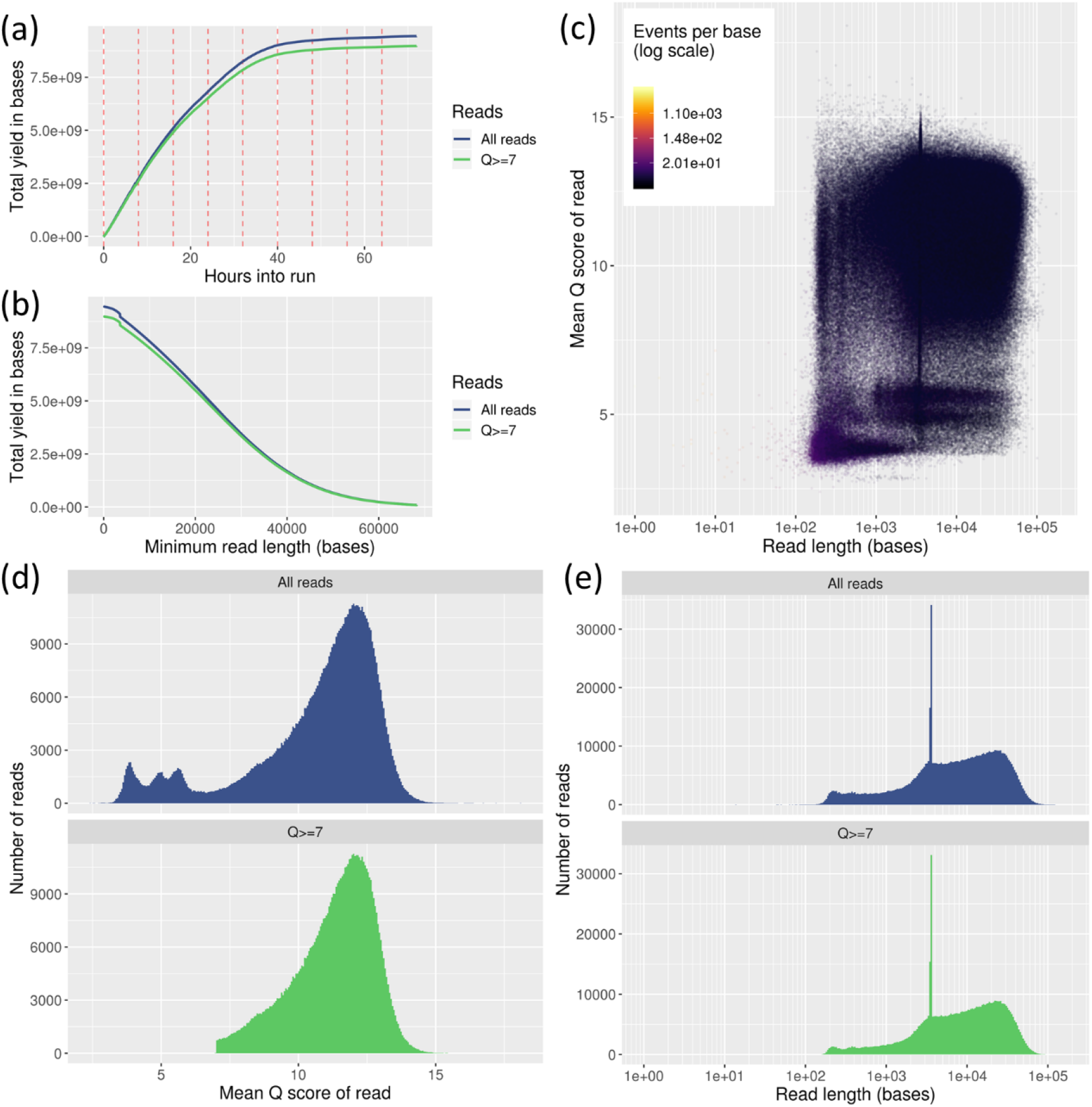
Long-read sequencing summary. (a) Yield by hour; (b) Yield by length; (c) Q score of bases and reads by read length; (d) Histogram of read length; (e) Histogram of Q score.

### 3.2 MinION and Illumina *de novo* genome assembly

A complete chromosome-level *de novo* genome assembly of *C. cinerea* strain #326 was obtained. Nanopore long reads were used to make scaffolds. Illumina short reads were used to correct the sequencing errors. Illumina re-sequencing generated 9.2 million reads and 1.38 Gb bases, with 98 % of bases had over 99 % accuracy. After quality filtering, a total of 8.97 Gb of long reads and 1.33 Gb of short reads were used to polish the strain #326 genome. The estimated coverage of long reads and short reads were 200 times and 35 times, respectively.

After cleansing and contamination removal, the assembly contained 31 contigs and all thirteen chromosomes of *C. cinerea* were assembled into single contigs, with the N50 of 2.8 Mbp. The thirteenth longest scaffold was 1,109,935 bp in length and about 10-fold longer than the following scaffolds. The longest thirteen scaffolds made up the assembly size of 37,536,632 bp (Table S2). Compared with the previously released Illumina short reads genome assembly of strain #326, our hybrid assembly reduced the number of contigs by 30-fold and increased N50 by 28-fold (Table 1).

**Table 1.** Comparison of three genome assemblies of *C. cinerea.*

| Strain | *A43mut B43mut pab1-1* #326 |  | Okayama-7 #130 |
| --- | --- | --- | --- |
| Source | Illumina<br>(Muraguchi et al., 2015) | Nanopore + Illumina<br>(This study) | Whole genome shotgun sequencing<br>(Stajich et al., 2010) |
| **Contigs (> 1 Kb)** | 944 | 31 | 68 |
| **Largest contig (bp)** | 622,701 | 4,326,744 | 4,146,986 |
| **Smallest contig (bp)** | 1,000 | 7,943 | 2,208 |
| **N50** | 100,685 | 2,822,236 | 3,468,139 |
| **Total assembly size** | 34,118,803 | 38,741,595 | 36,192,590 |
| **GC content (%)** | 51.62 | 51.57 | 51.64 |
| **BUSCO (basidiomycota_odb10, n=1764)** | 1737 (98.5%) | 1735 (98.4%) | 1735 (98.4%) |
| Complete and single copy | 1729 (98.0%) | 1721 (97.6%) | 1730 (98.1%) |
| Complete and duplicated | 8 (0.5%) | 14 (0.8%) | 5 (0.3%) |
| Fragmented | 10 (0.6%) | 11 (0.6%) | 9 (0.5%) |
| Missing | 17 (0.9%) | 18 (1.0%) | 20 (1.1%) |

Manual curation resolved the mitochondrial genome. In the first draft assembly, the mitochondrial scaffold of was twice the length of Okayama-7 #130 mitochondrial genome (85,870 bp versus 42,448 bp). The mitochondrial scaffold of strain #326 went around the mitochondrial genome twice. This problem is a common artefact when assembling circular genomes using Canu (Koren et al., 2017). Reconstructing mitochondrial genome using circular DNA assembler, Unicycler (Wick et al., 2017), following the hybrid assembly method resolved the problem. The complete mitochondrial DNA sequence was 42,417 bp in length, with equivalent quality as the linear scaffolds. In short, the final *C. cinerea A43mut B43mut pab1-1* #326 assembly contains 13 chromosome-level scaffolds and one complete mitochondrial genome (Table S2). Sequencing raw reads could be aligned to the genome assembly with a mapping rate of 94.39 % and 99.21 % for long reads and short reads, respectively.

### 3.3 Completeness of the genome

Genes recovered from assembly evaluated using BUSCO with basidiomycota_db10 database and showed a 98.4 % degree of completeness. A total of 1,735 complete basidiomycota core genes were found with 1,721 single copy and 14 duplicated genes (Table 1). There were more duplicated genes than previous genome assemblies. These duplications can be artefacts from scaffolding, where error-prone long reads were used (Minei et al., 2018). However, long reads take advantages in resolving repetitive regions, which could help distinguishing homologues and increase the number of duplicated core genes (Díaz-Viraqué et al., 2019). In addition, 11 core genes were fragmented, and 18 genes were missing in the assembly.

The draft genome assembly was compared with strain #130 reference genome with several methods. In strain #130 reference genome, the longest 13 contigs were designated as 13 chromosomes, with an assembly size of 35,810,191 bp (Stajich et al., 2010). Sequence alignment and blast search showed that most genomic regions in strain #326 have over 99 % sequence similarities against strain #130 (Fig. S2). Seventy-two regions displayed alignment lengths of over 100 Kbp against strain #130. The longest 13 scaffolds corresponded to 13 chromosomes in strain #130 genome with no chromosomal fusion or translocation. Matching to the strain #130 reference genome, seven chromosome-level scaffolds of strain #326 genome assembly were forward-oriented, and six scaffolds were reverse-oriented. Genome sequences of these strains had high homology (Fig. 4a). All-vs-all comparison showed that 96.33 % of the bases in strain #130 genome and 95.71 % in #326 genome aligned to the cross references. Sequences of 34.7 Mbp in 1,608 alignments were orthologs, with 99.11 % average identity. MCscan identified 12,062 orthologous gene pairs in the coding sequence annotation of these genomes. The gene pairs could be grouped into 55 clusters. A simplified karyotype plot revealed the macrosynteny of two genomes (Fig. 4b). Analysis results based on genome sequences and coding regions were consistent.

**Fig. 4.**
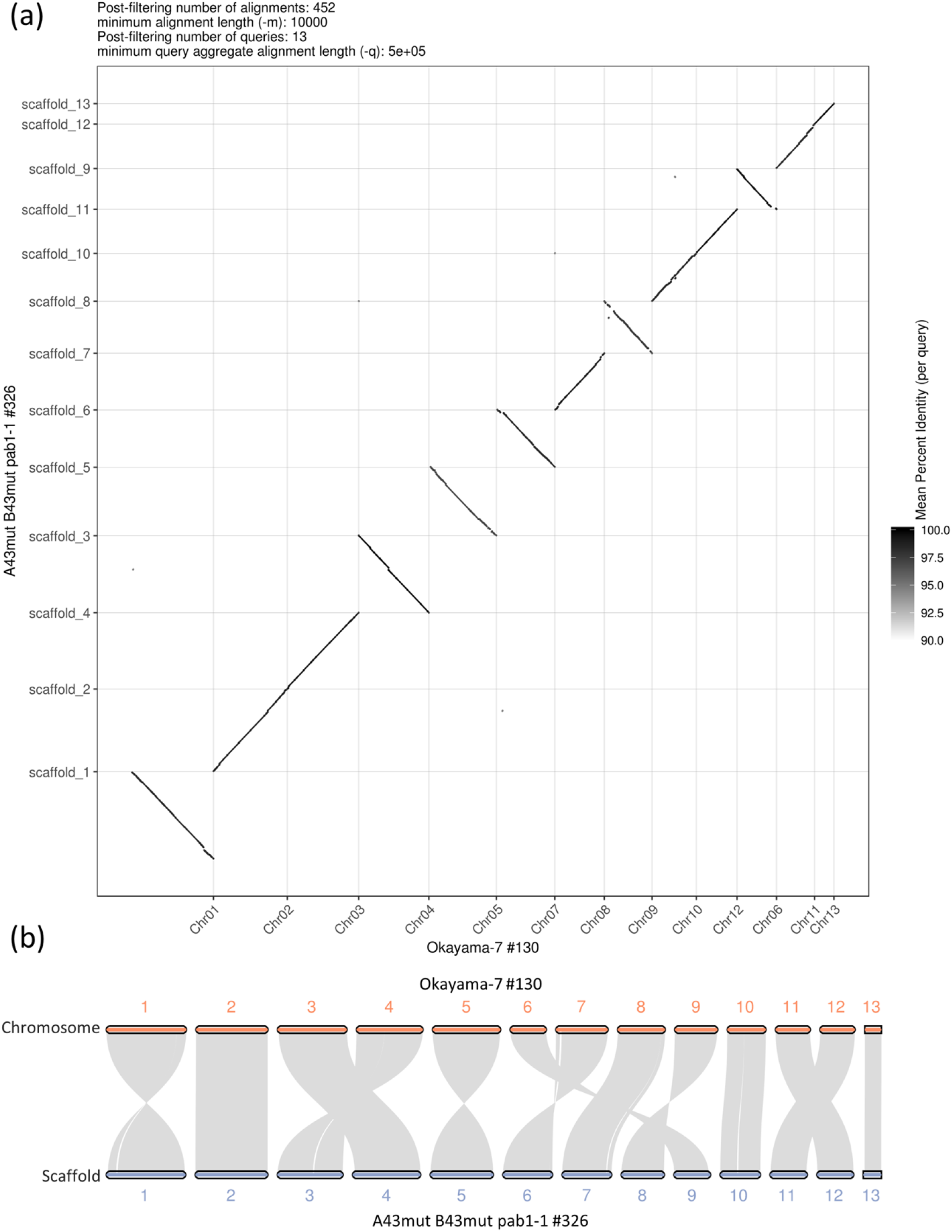
Genome comparisons between two *C. cinerea* strains. (a) Draft genome alignment result generated by MUMmer package, 1-to-1 alignment with minimum alignment length set to 10 Kbp. (b) Simplified karyotype plot, showing coding sequence based macrosyntheny of genomes.

We also tested the relationship between sequencing depth and genome sequence contiguity using randomly sampled ONT long reads. Nanopore sequencing depth below 120-fold may split a chromosome into several scaffolds (Fig. S3). With 30-fold sequencing depth, split and fusion of chromosomes were observed in the genome assembly. At least five of thirteen chromosomes were fragmented and assembled into over 11 scaffolds. Increasing the sequencing depth to 60-fold solved the problem. Two of three trials displayed that only chromosome II was split into two scaffolds, and the other 12 chromosomes were completely well assembled. However, the fragmentation problem of the chromosome II remained when the subsequence coverage was increased to 90-fold. All three assembly trials with 120-fold sequence coverage gave the chromosome-level assembly of all 13 chromosomes.

### 3.4 Gene prediction and functional annotation

Eighteen transcriptomes, including developmental stages of vegetative mycelium, oidiation, sclerotia formation, hyphal knots, primordia and young fruiting bodies (Xie et al., 2020), were used to support the genome annotation. A total of 15,250 genes was predicted in the genome, with 16,862 transcripts. Table 2 summarised the gene annotation results in three assemblies of *C. cinerea.* Compared with the previous genomes, there is an increase in gene number and average gene length in the hybrid genome assembly. The total gene length was 2.5 Mbp longer than in short-read constructed genome. The total number and total length of coding sequences, exons and introns increased in the hybrid genome assembly, but the average length was similar to other assemblies. For the untranslated regions, more 5’-untranslated regions (UTR) but less 3’-UTR was annotated in hybrid genome assembly than in Illumina short reads-constructed genome. It also showed that in strain #326, the number of 5’-UTR was about two times more than in strain #130, while the number of 3’-UTR was only half of strain #130. Although the number of regions varied, the length of UTR was significantly longer in strain #326, especially in the hybrid genome assembly.

**Table 2.** Comparison of gene annotation results in three assemblies of *C. cinerea.*

| Strain | *A43mut B43mut pab1-1* #326 |  |  |  |  |  |  |  | Okayama-7 #130 |  |  |  |
| --- | --- | --- | --- | --- | --- | --- | --- | --- | --- | --- | --- | --- |
|  | Nanopore + Illumina<br>(This study) |  |  |  | Illumina<br>(Muraguchi et al., 2015) |  |  |  | Whole genome shotgun sequencing<br>(Stajich et al., 2010) |  |  |  |
| Source | Number | Average<br>Number | Length | Average<br>Length | Number | Average<br>Number | Length | Average<br>Length | Number | Average<br>Number | Length | Average<br>Length |
| **Gene** | 15,250 | / | 27,402,013 | 1,796.9 | 14,242 | / | 24,898,399 | 1,748.2 | 13,393 | / | 23,611,982 | 1,763.0 |
| **CDS** | 80,939 | 5.3 | 19,540,500 | 1,281.3 | 75,485 | 5.3 | 18,519,347 | 1,300.3 | 75,320 | 5.6 | 18,522,018 | 1,383.0 |
| **Exon** | 83,580 | 5.5 | 22,650,030 | 1,485.2 | 77,000 | 5.4 | 20,450,942 | 1,436.0 | 75,796 | 5.7 | 19,109,420 | 1,426.8 |
| **Intron** | 68,330 | 4.5 | 4,751,983 | 311.6 | 62,758 | 4.4 | 4,447,457 | 312.3 | 62,403 | 4.7 | 4,502,562 | 336.2 |
| **5'-UTR** | 7,096 | 0.47 | 1,410,022 | 92.5 | 6,725 | 0.47 | 885,532 | 62.2 | 2,409 | 0.17 | 185,417 | 13.9 |
| **3'-UTR** | 5,945 | 0.39 | 1,679,083 | 110.1 | 6,260 | 0.44 | 1,046,063 | 73.4 | 13,522 | 1.0 | 401,985 | 30.0 |

Further analysis functionally annotated 11,704 genes, which is 76.7 % of the total gene count (Table 3). Among the protein sequences, 90.6 % matched strain #130 sequence records (taxid: 240176, e-value < 1e-5). Based on the BLAST search, 10,274 protein sequences were assigned with Gene Ontology terms. A total of 7,356 gene sequences got Pfam annotations. Eggnog-mapper v5.0 (Huerta-Cepas et al., 2018) annotated 6,547 genes with Cluster of Orthologous Groups (COG). Based on the protein sequences, 1,796 signal peptides and 2,918 transmembrane proteins were annotated using Phobius (Käll et al., 2004). In addition, 342 peptidases and 449 CAZymes were annotated by merops (Rawlings et al., 2018) and dbCAN (Zhang et al., 2018), respectively. tRNAscan-SE (Lowe and Eddy, 1996) annotated 238 tRNA coding genes in the assembly, which was 53 genes less than in the strain #130 genome annotation. Repetitive contents took up about 2 % of the genome sequence (Table 4).

**Table 3.** Functional annotation summary on the novel assembly.

| **Annotation** | **No. of genes** |
| --- | --- |
| **GO** | 10,274 |
| **Pfam** | 7,356 |
| **COG** | 6,547 |
| **Signal peptides** | 1,796 |
| **Transmembrane proteins** | 2,918 |
| **Peptidases** | 342 |
| **CAZyme** | 449 |
| **tRNA coding** | 238 |

**Table 4.** Statistic of repetitive contents in three assemblies of *C. cinerea.* Long terminal repeats (LTR) are retrotransposons, non-LTR retrotransposons comprise short interspersed nuclear elements (SINE) and long interspersed nuclear elements (LINE).

| Strain | *A43mut B43mut pab1-1* #326 |  |  |  | Okayama-7 #130 |  |
| --- | --- | --- | --- | --- | --- | --- |
| Source | Nanopore + Illumina |  | Illumina |  | Whole genome shotgun sequencing |  |
|  | (This study) |  | (Muraguchi et al., 2015) |  | (Stajich et al., 2010) |  |
|  | Number of elements | Length occupied (bp) | Number of elements | Length occupied (bp) | Number of elements | Length occupied (bp) |
| **Interspersed** |  | 18,668 |  | 9,804 |  | 12,530 |
| SINE | 34 | 2,130 | 22 | 1,287 | 30 | 1,764 |
| LINE | 110 | 8,366 | 88 | 6,748 | 106 | 8,174 |
| LTR | 12 | 1,368 | 3 | 204 | 12 | 1,383 |
| DNA | 25 | 2,019 | 21 | 1,565 | 15 | 1,117 |
| Unclassified | 25 | 4,785 | 0 | 0 | 2 | 92 |
| **Simple repeats** | 6,239 | 279,133 | 5,365 | 236,661 | 5,786 | 258,099 |
| **Low complexity** | 1,173 | 63,623 | 959 | 51,013 | 1,150 | 62,318 |
| **Small RNA** | 755 | 465,192 | 125 | 8,718 | 130 | 12,445 |

The hybrid genome assembly of strain #326 was about 4.6 Mb larger than the short-read assembly. To identify the differences, the scaffolds in short-read constructed genome was aligned to the hybrid assembly using minimap2 v2.17 (Li, 2018). The new sequences spread out over the genome (Fig. S4a). Within the 4.6 Mbp unaligned regions, there were 879 genes (1,125,473 bp) and about 530 Kbp repeated regions, which took up 65 % of the repeated regions found in the whole genome. 856 Kbp unaligned sequences were located at the terminals of the scaffolds. In addition, among those 755 small RNAs found in the whole genome, 727 of them were in the unaligned region. By pairwise blast search against the protein coding sequences annotated in three genome assemblies, a total of 11,503 genes were found common to all genomes, and 2,602 genes were found specific in the hybrid genome assembly (Fig. S4b).

### 3.5 Single nucleotide variant identification

The genome assembly and annotation were applied to downstream studies. Six selected mutants were sequenced on MinION platform, including three fruiting mutants (UV423, UV506 and UV512) and three non-fruiting mutants (UV433, UV554 and UV663). The morphology of the mutants was shown in Fig. S5. The use of high contiguity reference genome improved the mapping rate by 2-5 % (Table S3). Aligning mutant sequences to the reference genome, 496 SNVs were identified in these six mutants, locating in 441 genes or intergenic regions (dataset S2). On the other hand, aligning the mutant sequences to the short reads-constructed genome, there are ten times more candidate sites (Table S3), with 75 % SNVs occurred in more than one mutant. The high contiguity genome improves alignment accuracy and cuts down the systemic error in SNV calling.

Referring to the hybrid genome assembly and annotation, around 40 % of the SNVs were found in coding regions. Point mutations in three hypothetical proteins, CC2G_003551, CC2G_013038 and CC2G_015270, were found in more than one isolates. The C-to-T transition at DNA mismatch repair protein *MSH6* (CC2G_OI3873 c.l483C>T, scaffold_8:lO37634C>T) was validated by Sanger sequencing in non-fruiting isolate UV663 (Fig. S6a). One CAA codon, encoding Gln, was changed into TAA, creating a stop codon that caused an early termination of the protein. *MSH6* was differentially expressed during fruiting body development of strain #326 (Fig. S7). The expression level greatly increased in primordia and young fruiting body. In UV663, expression of the gene was significantly down-regulated during vegetative growth, and slightly increased during hyphal knot formation (Fig. S7). UV663 failed to form fruiting body, and developed clustered fruiting body initials instead (Fig. S2g). In another non-fruiting isolate UV433, the G-to-A transition at atypical/alpha protein kinase (CC2G_008473 c.55+1G>A, scaffold_4:5425l5G>A) was identified and validated (Fig. S6b). It is predicted to change the splicing site of the transcript. However, the expression level of the gene was very low and undetectable by RT-qPCR.

### 3.6 Structural variant identification

To better determine the minimum coverage or number of supporting reads for our data, we navigated a bunch of sequenced mutants with the average coverage varied from 2.89 × to 25.11 ×. The higher requirement on number of SV supporting reads, the less variants were called (Fig. S8). When average coverage is higher than 10-fold, there is a flat-stage on SV number at four to six supporting reads. The minimum requirement of five supporting reads was finalised, considering both accuracy and the uneven distribution of sequencing depths across the genome.

With the criteria of at least 5 supporting reads, 2 insertions and 3 deletions were called in six mutants in reference to our genome assembly (dataset S3). We tried performing SV calling using short read-constructed reference genome with the same criteria, and 766-1016 SVs were found in specific mutant (Table S3). Over 90 % of these SVs were notes as breakpoints. Manual curation found that the two ends of these SVs located at the start point or end point of two different scaffolds. Those scaffolds were later proved to be continued on the chromosomes. The chromosome-level reference genome gets rid of the scaffold breakpoint problems and improves the prediction accuracy. Therefore, a high contiguity reference genome is important for long read sequencing data analysis.

Referring to our genome assembly, a 66 bp deletion on Ras GTPase-activating protein *(RasGAP,* CC2G_009974) was found in all mutants. This deletion was also identified when referring to the short read-constructed strain #326 reference genome. The *RasGAP* deletion in six selected mutants were validated by Sanger sequencing (Fig. S9a). Other screened mutants were also tested by PCR, and gel electrophoresis results showed that all 100 mutants we collected have the same deletion on *RasGAP* gene (Fig. S9b). RNA of the 6 sequenced mutants was isolated from 4 days vegetative mycelia and 2 days hyphal knot inducing mycelia, and the expression of *RasGAP* can be detected. Sanger sequencing on reverse transcribed cDNA showed that the deletion changes the splice site of the transcript and would result in the early termination of the protein sequence (Fig. 5). RT-qPCR results showed that *RasGAP* (CC2G_009974) was not differentially expressed during the transition from vegetative growth to hyphal knot formation in strain #326 wildtype. However, in non-fruiting isolates UV554 and UV663, the gene expression level was significantly up-regulated during development (Fig. S10). The 66 bp deletion in CC2G_009974 was not located at the *RasGAP* functional domain.

**Fig. 5.**
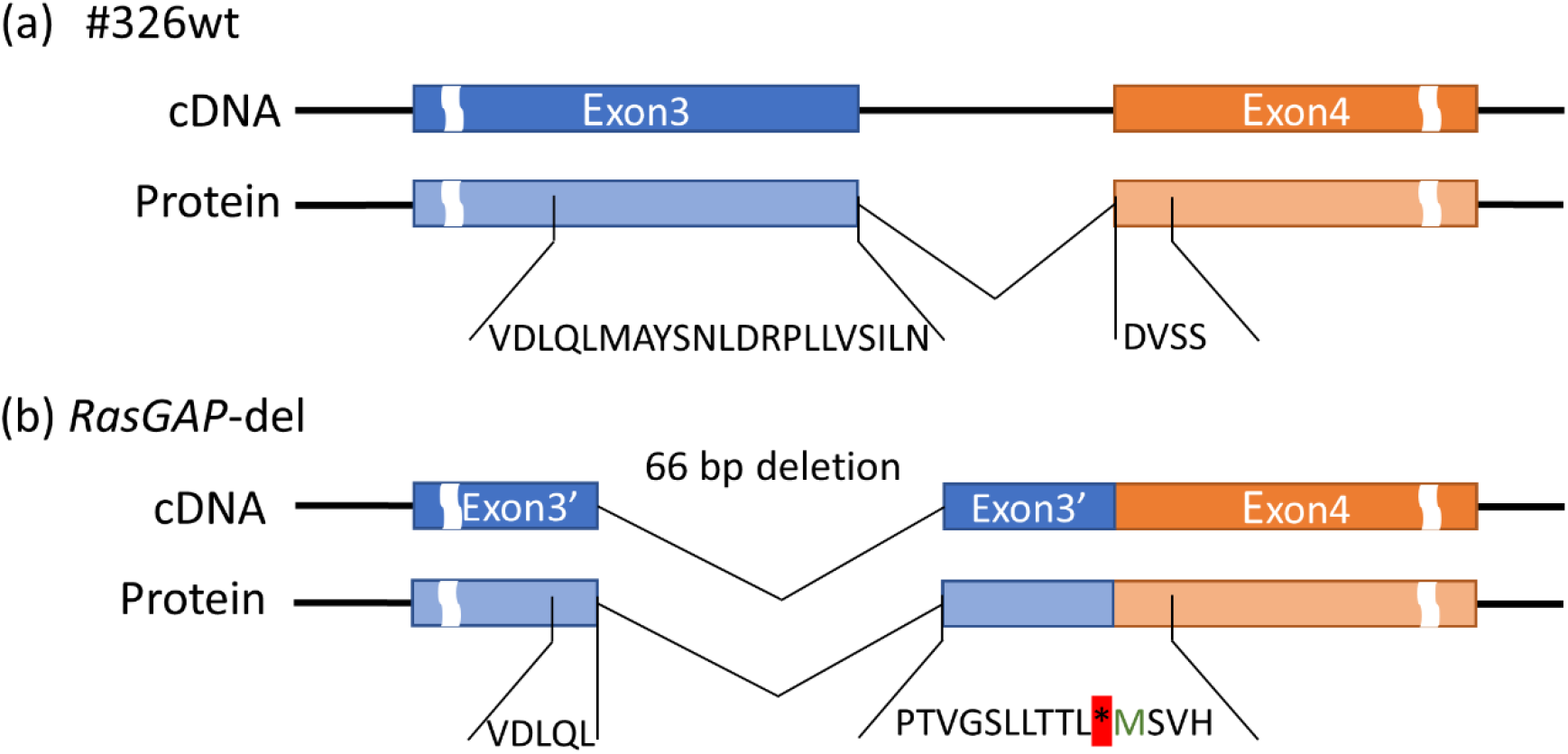
The 66 bp deletion in *RasGAP.* (a) cDNA and protein sequence of *RasGAP* in #326 wildtype. (b) cDNA and protein sequence of *RasGAP* in mutants. Sequence validation by Sanger sequencing are shown in Fig. S8.

## 4. Discussion

In this study, we presented a series of fungal genomic analyses based on MinION sequencing platform. We modified an efficient HMW gDNA isolation protocol, and that is suitable for fungal genome sequencing with MinION. We have constructed a near-complete chromosome-level genome assembly of *Coprinopsis cinerea* strain *A43mut B43mut pab1-1* #326. All 13 chromosomes and the mitochondrial genome of *C. cinerea* were assembled into single scaffolds. This assembly is unfragmented and has high degree of synteny with *C. cinerea* Okayama-7 #130 reference genome. In addition, we applied the assembly to identify structural variants and single nucleotide variants in mutants of strain #326. Our results demonstrated the possibility of high-quality assembly of a middle-sized fungal genome and fast screening of genotype of fungal isolates using MinION sequencing platform. An optimal series of experimental and *in silico* references for fungal genomic analyses were provided.

### 4.1 Magnetic bead based HMW gDNA extraction is sufficient for high quality genome sequencing and assembly of *C. cinerea*

In this study, we modified a high-performance and cost-effective HMW genomic DNA extraction methods for fungal samples. Due to the complexity of multi-layered cell walls and high polysaccharides and pigments concentrations, isolating high quality DNA from fungal samples is challenging (Lee et al., 2017). Compared with CTAB or phenol-chloroform methods (Fauchery et al., 2018; Gong et al., 2019), the use of magnetic beads gets rid of hazardous organic solvents and their contamination problem. The magnetic bead protocol has low requirements on laboratory setup and experimental skills. Our results demonstrated that such method offers convenience to ONT sequencing experiments and other studies that have requirements on long DNA fragment, especially in fungal samples with high polysaccharide content. It can be further adapted to a broad range of samples by changing the cell lysis condition. The unit price of the sample is relatively low while the yield is high. Although the proportion of ultra-long reads (read length > 50 Kbp) was not as high as expected in our sequencing data, it gave valuable data to the genome assembly. Our assembly results showed that long DNA fragments, but not ultra-long DNA fragments, is enough to obtain the high-quality assembly of a fungal genome. Therefore, megabase-sized DNA obtained by plug extraction would be a good addition but not a necessity for fungal genome construction (Jain et al., 2018). Our results proved that DNA sample isolated using magnetic beads is quality enough for MinION library preparation and sufficient for the construction of a middle size genome.

### 4.2 Feasibility of fungal genomic studies using ONT

With the use of Nanopore long reads, we obtained the chromosome-level genome sequence of *C. cinerea* strain #326. Long reads generated from third generation sequencing platforms improve contiguity and completeness of genome assemblies. Although the assembly size of long reads and short reads remains largely equal in some cases, sequence addition and deletion are both common in long read assemblies (Dutreux et al., 2018; Johnson et al., 2020). Our hybrid assembly showed 28-fold decrease in number of contigs, with 4.6 Mb increase in assembly size (~12 %). Similar to the observation in *Saccharomyces cerevisiae* (Salazar et al., 2017), these additional sequences contain many protein coding genes. In the long read assembly of *Trypanosoma cruzi,* the assembly size increased 60 % and a gene recovery increased 3-fold, especially in the identification of repetitive coding regions (Díaz-Viraqué et al., 2019). The hybrid assembly of *C. cinerea* strain #326 also showed a 2-fold increase of interspersed contents, which was the same as the case of *Parochlus steinenii* (Shin et al., 2019). The use of long reads helps resolving homologues and repetitive regions in the genome (Bongartz, 2019; Jain et al., 2018).

Compared to next generation sequencing technologies, ONT reads have the problem of relative low accuracy. A high quality *de novo* genome assembly using only ONT reads would require high sequencing quality on base accuracy, long read length, and great sequencing depth (Tyson et al., 2018; Vaser et al., 2017), increasing the sequencing cost. By introducing the high accuracy short reads to the assembly polishing, most errors could be corrected as long as the long reads error rate was below 15 % (Wang and Au, 2020), and the requirement on long read coverage can be decreased. The improvement on sequence accuracy provides great advantages to the BUSCO score and gene annotation (Johnson et al., 2020). Our results demonstrated that ONT long read with 60-fold coverage and short read polishing could yield a high-quality chromosome-level *C. cinerea* genome. Such hybrid genome assembly approach lowers the threshold of ONT read coverage and sequencing time and cost to about a quarter of the long read only assembly approach.

Fungi are a group of diverse organisms with estimated 2.2 to 3.8 million species (Hawksworth and Lücking, 2017). Among the 120,000 well described fungal species, there are great differences in all aspects, including structural and physiological features, living forms, lifestyles and life cycles (Halbwachs et al., 2016; Nagy et al., 2017; Treseder and Lennon, 2015). Despite these differences, most fungal species exist as hyphae. Hyphae are long and branched filamentous structures made up of thread-like and end-to-end connected cells, with the cell wall composed of chitin, glucans and glycoproteins. Hyphae are the main form of vegetative growth and the basic unit of well-organized fruiting body structures (Bowman and Free, 2006; Krizsán et al., 2019; Nguyen et al., 2017). On the other hand, although the genome sizes of fungi varies by almost three orders, a large proportion of fungal species has the genome size of 20 ~ 60 Mbp, average around 40 Mbp (Stajich, 2017). Thus, the throughput of a MinION flowcell (8 ~ 10 Gb in 72 h) is sufficient and cost-effective to fungal genomes (Díaz-Viraqué et al., 2019; Lee et al., 2019). Our workflow should be applicable to other fungal genome analyses because fungi have similar cellular structures and genomic features.

### 4.3 ONT is applicable in genome resequencing and genetic variant detection

Genome resequencing with ONT platforms have been performed in many studies of microbes, plants, animals and human (Bowden et al., 2019; Michael et al., 2018; Miller et al., 2018; Salazar et al., 2017). The long reads make it easy to obtain contiguous genome sequence, to resolve complex genome regions, and to detect structural variants. In this study, we resequenced six mutants, and identified genetic variants by comparing our strain #326 reference genome. We compared the read alignment and variant calling using two different reference genomes, the hybrid genome assembly and the short read based genome assembly. The alignment rate improved by 2-5 % with the high contiguity reference genome. However, the numbers of SV and SNV found from fragmented reference genome was 100-times and 10-times more than the chromosome-level genome (Table S3). Most of the SVs and SNVs found with fragmented reference genome were false positive events caused by misalignment and scaffold breakpoints. They raised difficulties in variant validation and mutant genotyping. A high contiguity reference genome is essential to genetic and genomic studies. To date, several tools and pipelines have been developed to call SVs in the genome (Zhou et al., 2019). Previous studies suggested a minimum average coverage of 10-fold to ensure a comprehensive detection of SV using nanopore reads (Cretu Stancu et al., 2017; De Coster et al., 2019; Gong et al., 2018). Our navigation on relationship between identified SV number and average coverage/supporting read number agreed with those reports. To make it one step further, we also look into the read alignment results of low sequencing coverage isolates (< 5 ×). The *in silico* prediction of SV sites were reasonably accurate in the mutants. The read alignment and blast against the genome confirmed that the parameters had sufficiently called SV from the reads and filtered most false-positive events. The results showed the potential of fast screening of SVs with low sequencing coverage, which can be useful in off-target identification of CRISPR systems (Zischewski et al., 2017) and many strain selection systems (Gowers et al., 2020).

Due to the relatively high error rate, ONT reads were suggested not to be used in single nucleotide variant calling, especially when the average coverage was below 20 × (Magi et al., 2017). Recently, the base accuracy of ONT reads has been greatly improved by the better sequencing chemistry and basecalling algorithm. To date, SNV calling from ONT reads can be achieved by assembly-based and alignment-based approaches (Guo et al., 2018; Loman et al., 2015; Malmberg et al., 2019). The application of the neural network also improves the accuracy but is costly due to intensive computations (Luo et al., 2019). Here, we performed SNV calling using “minimap2+bcftools” method, with the restriction of at least 5 supporting reads and the alternative ratio over 90 %. Considering the average base accuracy of ~ 90 %, these parameters would greatly filter out the sequencing errors and potential mixture of cell isolates (*P* < 1 × 10^-4^). However, systemic errors from basecalling and alignment were not fully excluded. Further filtration and validation, as well as purification and backcross are needed for phenotyping and functional characterisation of the isolates (Zuryn et al., 2010).

### 4.4 Potential impacts of *RasGAP* deletion

We have found a common 66 bp deletion on *RasGAP* in the selected mutants. *Ras* proteins are important regulators of signal transduction in cells. It is involved in the regulation of several cellular processes through the RAS/cAMP/PKA pathway, including cell cycle, meiosis and autophagy (Huang et al., 2019; Repasky et al., 2010). The catalytic activities of Ras proteins are regulated by RasGAP and Ras guanine nucleotide exchange factors (GEFs). RasGAP converts GTP-bound Ras to GDP-bound Ras and inactivate the Ras protein, and GEF performs the reaction in the reverse direction (Tamanoi, 2011). *RasGAP* regulates the light responses and circadian clock in *Neurospora crassa* (Polaino et al., 2017). In *Magnaporthe oryzae, RasGAP* is required for conidial morphogenesis and plant infection (Kershaw et al., 2019). In *Schizophyllum commune, RasGAP* regulates hyphal growth orientation and fruiting body development (Schubert et al., 2006). However, we have not observed similar changes in the mutants except the decrease in growth rates. A total of thirteen *Ras* proteins and two *RasGAP* (CC2G_007150 and CC2G_009974) have been annotated in this *C. cinerea* assembly. Further, the AGC/PKA protein kinase (CC2G_008645) and CMGC/MAPK protein kinase (or glycogen synthase kinase, CC2G_005966) are two of the potential downstream kinases of *RasGAP* (Yamamoto-Honda et al., 1995). Other genes related to the regulatory pathways, including adenylate cyclase, glycogen synthase, glycogen phosphorylase, trehalase and trehalose phosphate, were taken into consideration as well (Santangelo, 2006; Thammahong et al., 2017). We observed that the gene expression levels of these potential downstream targets in the pathway varied in different isolates (data not shown). We referred the inconsistency and unpredictability of phenotypes to the wide-spread background mutations and the complexity of regulatory network (Chandler et al., 2013; Leberer et al., 2001). Although the deletion was not in the functional domain of *RasGAP,* this deletion looks similar to the cleavage of RasGAP peptide in HCT116 cell lines under the mild stress induced by cisplatin. Such cleavage is considered to be essential for cell survival under stress (Yang et al., 2004; Zhang et al., 2014, 2011). The similarity and difference on regulation of the two *RasGAP* genes and the deletion impacts on the fungus remains largely unknown.

## 5. Conclusion

Our results provided a series of approaches and reference parameters for fungal genomic analysis. Our improved procedure includes DNA isolation, whole genome sequencing, genome assembly and annotation, and mutant genotyping. We have constructed a chromosome-level genome assembly of *C. cinerea A43mut B43mut pab1-1* #326, and identified the genetic variants in #326 mutants. This workflow is flexible and expandable, and would be applicable to different fungal studies.

## Acknowledgement

We would like to thank Dr. Hajime Muraguchi for kindly sharing the *C. cinerea* strain #326 with us. We would also like to thank Dr. Chun Hang Au for fruitful discussions and Ms. Lei Xing for help on experiments.

